# Somatic mutations in ALS genes in the motor cortex of sporadic ALS patients

**DOI:** 10.1101/2025.03.31.646375

**Authors:** Óscar González-Velasco, Rosanna Parlato, Rüstem Yilmaz, Lorena Decker, Sonja Menge, Axel Freischmidt, Xiaoxu Yang, Nikshitha Tulasi, David Brenner, Peter M. Andersen, Karin M.E. Forsberg, Johannes C.M. Schlachetzki, Benedikt Brors, Lena Voith von Voithenberg, Jochen H. Weishaupt

## Abstract

Amyotrophic lateral sclerosis (ALS) is characterized by the progressive degeneration of cortical and spinal motor neurons. Mendelian germline mutations often cause familial ALS (fALS) but only approximately ten percent of sporadic ALS cases (sALS). We leveraged DNA and single cell RNA-sequencing data from autopsy tissue to explore the presence of somatic mosaic variants in sALS cases. Deep targeted panel sequencing of known ALS disease genes in motor cortex tissue revealed an enrichment of low allele frequency variants in sALS, but not in fALS with an identified monogenic cause. *In silico* analysis predicted increased pathogenicity of mosaic mutations in various known ALS mutational hot spots. In particular, we identified the somatic FUS variant p.E516X, located in an established hot spot for germline ALS mutations, which leads to nucleo-cytoplasmic mislocalization and aggregation typical for ALS FUS pathology. Additionally, we performed somatic variant calling on single cell RNA-sequencing data from sALS tissue and revealed a specific accumulation of somatic variants in excitatory neurons, reinforcing a neuron-autonomous disease initiation. Collectively, this study indicates that somatic mutations within the motor cortex, especially in excitatory neurons, may contribute to sALS development.

## Introduction

Amyotrophic lateral sclerosis (ALS) is an adult-onset neurodegenerative disease characterized clinically by degeneration primarily of motor neurons eventually leading to respiratory failure and death^1,2^. Only about 5-10% of European ALS patients report a positive family history for the disease (fALS)^3^, usually with an autosomal dominant mode of inheritance, whereas most ALS patients report a family history that is unremarkable for ALS (sALS). Rare cases of germline *de novo* mutations, in particular in *FUS* and *SOD1* as a cause of sALS, have been reported^4,5^. However, whereas even in a considerable proportion (∼50%) of fALS cases screening for a germline mutation in blood DNA remains inconclusive, conventional genetic testing turns out negative in approximately 90% of the people with sALS^6^. On the other hand, twin studies point to a considerably higher contribution of genetic factors to ALS pathogenesis than explained by the frequency of Mendelian mutations^7^.

Insights from neuropathology, but also progression of clinical symptoms, suggest that ALS pathology starts focally as a proteinopathy^8,9^ and then spreads contiguously within the central nervous system over time^10^. Biological and clinical data support the notion that a prion-like mechanism with focal initiation may be involved in spreading toxic RNA and protein species including misfolded SOD1, TDP43 and FUS^11,12^. Such a hypothesis entails that even a small number of pathologically altered cells may be sufficient to initiate the focal development of ALS pathology, with expanding motoneuronal demise, and eventually leading to clinically progressive manifest muscle weakness^13^. Studies in chimeric mice transgenic for fALS *SOD1* mutations suggest a neuron-autonomous initiation of ALS^14^. Furthermore, a combination of multiple genomic alterations may increase the risk of ALS development^15^. Considering these premises, here we tested the hypothesis that somatic mosaic mutations could account for a proportion of sALS of unknown monogenic origin, and aimed at the identification of the most burdened neuronal populations. To this end, we combined deep targeted sequencing of ALS-related genes, targeted amplicon sequencing, and single cell RNA-sequencing of post-mortem motor cortex tissue followed by a proof-of-concept pathogenic validation of selected mosaic mutations in the ALS gene *FUS*. This study shows that somatic mutations in ALS-related genes may play a role in sALS pathogenesis, and that within the motor cortex excitatory neurons are more prone to accumulate somatic mutations.

## Material and Methods

### Patient cohort

Fresh frozen autoptic human precentral gyrus and spinal cord tissues of donors with ALS and control donors were provided by the ALS Brain Bank at Umeå University in Sweden and the Netherlands Brain Bank (Supplementary Table 1, Supplementary Materials and Methods). Patients who donated tissue to the ALS Brain Bank at Umeå University and the Netherlands Brain Bank provided written informed consent for the molecular genetic research reported in this manuscript.

### Targeted deep sequencing

Genomic DNA was extracted and 50 ng of genomic DNA was used for library preparation and target enrichment (Supplementary Materials and Methods). The list of ALS genes targeted in this study is provided in Supplementary Table 2. Next-generation sequencing was performed at the Sequencing Core Facility of the German Cancer Research Center (Supplementary Materials and Methods). The samples were sequenced on a HiSeq 4000 instrument (Illumina) by paired-end sequencing of 100 bp targeting a coverage of ≥2000x. Per sample ∼10-20 Mio reads were obtained. A schematic overview of the analysis workflow is provided in Figure S1 (Supplementary Materials and Methods).

### Functional validation of variants

The coding sequence of human wildtype *FUS* (NM_004960.4) was cloned into BglII and KpnI sites of pCMV-Myc-N (Clontech Laboratories, Mountain View, CA, USA). Whereas FUS p.R495X was already available^16^, other variants were introduced by site-directed mutagenesis as described in the manual of the QuikChange II Site-Directed Mutagenesis Kit (Agilent Technologies, Santa Clara, CA, USA). Respective oligonucleotides for *FUS* mutagenesis are listed in Supplementary Table 3. All sequences were verified by Sanger sequencing.

HEK293 cells were transfected with plasmid DNA using calcium phosphate precipitation with minor modifications^17^. 24 h post transfection, immunocytochemistry was performed as recently described^16^ using primary mouse anti-myc (1:500, Cell Signaling Technology, 2276) and secondary donkey anti-mouse-647 antibodies (1:500, Thermo Fisher Scientific, A-31571). Images were acquired on a Zeiss LSM 980 confocal microscope and analyzed using Image J.

### SNV calling from scRNA-sequencing data

Fastq files from a public single cell RNA-sequencing dataset^18^ were processed using cellranger version 8.0.1, and human genome reference version GRCh38. We then used SComatic for somatic variant calling (Supplementary Materials and Methods), annotations of groups of cells were defined using the data’s original cell types. In total we processed around 850.000 cells (number of samples: fALS=5, fFTLD=5, control=15, sALS=13, sFTLD=12).

### Statistical analysis

Statistical analyses of variant occurrence between disease categories were performed using negative binomial generalized linear models (GLM) including disease category (sALS, fALS, control) as the main predictor and sex, age, origin of patient, and total coverage as covariates to control for potential confounding effects (Supplementary Materials and Methods).

### Data availability

The raw sequencing data will be made available at the European Genome-Phenome Archive (EGAS00001008104).

## Results

### Higher somatic variant burden in ALS-related genes in sALS patients identified by targeted deep sequencing

To detect germline and somatic mutations in sALS brains by deep sequencing, we analyzed motor cortex DNA of 9 individuals diagnosed as sALS patients, 5 familial ALS (fALS) cases, which carried germline mutations in fALS genes (*SOD1, TBK1, NEK1*, and *C9ORF72*), and 6 control cases (Supplementary Figure S1, Supplementary Table1). SNVs were detected by combining error-correction based on unique molecular identifiers and calling and integration from multiple variant callers (Supplementary Figure S2). To identify somatic mosaic mutations by targeted deep sequencing, we specifically selected variants with low variant allele frequency (VAF). The overall relative distribution of SNVs and small insertions and deletions in the different genomic regions, e.g. exonic, 3’UTR, 5’UTR, intergenic, was similar between controls, fALS, and sALS patients (Supplementary Figures S3-S5). Interestingly, we observed an increase in the total number of somatic variants in sALS patients (μ=93.7, σ=28.8) compared to controls (μ=45.8, σ=26.6, t-test p-value=0.015) (Fig. 1A, Supplementary Figure S3D). No significant difference was noted between fALS and controls (p-value=0.59). An analysis of the number of SNVs by allele frequency (AF) showed an overall increase in the number of SNVs in the genes of interest for AF between 1% and 35% in sALS cases, which was driven by somatic variants (Fig. 1B-C). Whereas the majority of the genes known to play a role in fALS showed an increase in the number of variants in sALS patients versus controls, we observed a specifically strong enrichment in the number of SNVs in *VAPB, MAPT, FUS, NEFH, CCNF, NEK1*, and *TBK1*, for some of which the number of somatic SNVs almost doubled (Fig. 1D, Supplementary Table 4, Supplementary Text). SNVs detected in these genes are positioned in different protein regions, e.g. α-helical chains of the proteins, which may impact protein function (Fig. 1E). In summary, we identified an overall increased somatic variant burden and several pathogenic somatic mutations in ALS-associated genes in sALS patients, indicating somatic mosaic variants as potential contributors to sALS development.

**Figure 1.**
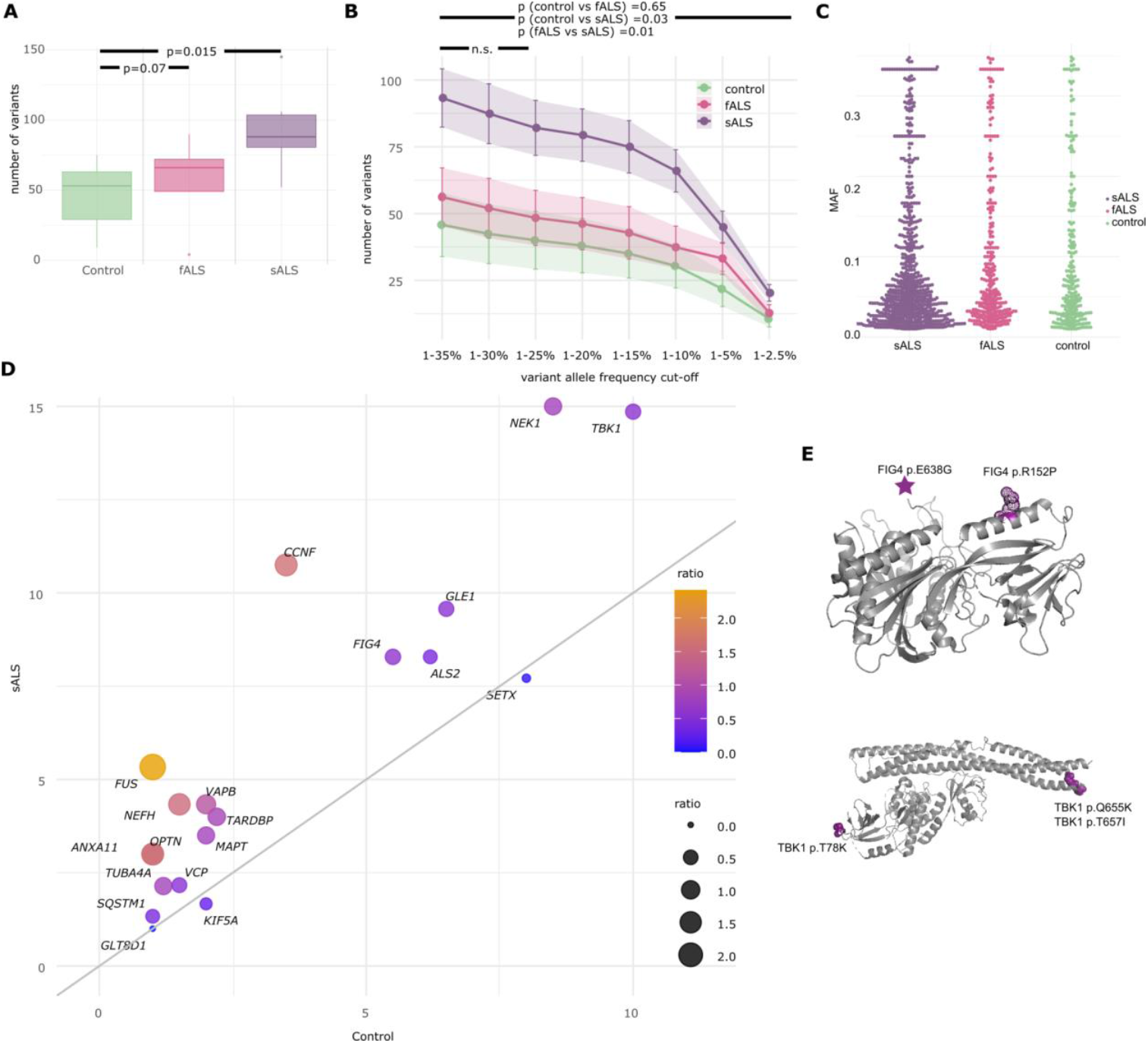
Somatic variant identification by targeted deep sequencing. (**A**) Number of variants with an allele frequency 35% > AF > 1% detected per sample in all genes of the targeted panel. (**B**) Number of variants detected per variant allele frequency for the different groups of samples. Enrichment test by using a generalized linear model with covariates (total coverage, age, sex, origin) and adjusted for multiple testing by Bonferroni approach. p values are provided for testing an allele frequency range of 0.35 < AF < 0.01. p values were non-significant (n.s.) for the allele frequency range 0.35 < AF < 0.25 for any of the conditions compared. (**C**) Allele frequency distribution of variants by group of samples. (**D**) Frequency of number of variants per sample in the indicated genes of the targeted panel in the control group versus the sALS group colored by the ratio of the number of variants between sALS and control samples. (**E**) Exemplary protein positions of SNVs detected in ALS samples displayed on the protein structures of FIG4 (pdb: 7K1W) and TBK1(pdb: 6NT9).

### Functional validation of somatic mosaic variants in FUS detected in sALS patients

ALS patients with a mutation in FUS show neuronal cytoplasmic aggregation of this protein^19,20^. To functionally validate the variants predicted to occur in a somatic mosaic pattern in FUS (Fig. 2A), we analyzed the cellular localization of the protein in cultured cells (Fig. 2B). Mutant FUS proteins carrying the 4 variants were cloned and transiently overexpressed in HEK293 cells in comparison to wildtype controls and a known variant (FUS p.R495X). Whereas wildtype FUS was localized in the nuclei in a well-defined pattern, the ALS-related FUS p.R495X variant showed cytoplasmic aggregation (Fig. 2B, C). The FUS p.E516X variant, which we detected in ALS patients in a somatic mosaic pattern, was also localized in aggregated foci in the cytoplasm, losing its nuclear localization. The cellular localization of FUS p.E516K, FUS p.E516E, and FUS p.G515V was only minimally affected under these experimental conditions, which is in line with pathogenic effects of FUS variants even in the absence of cytoplasmic aggregation^21^.

**Figure 2.**
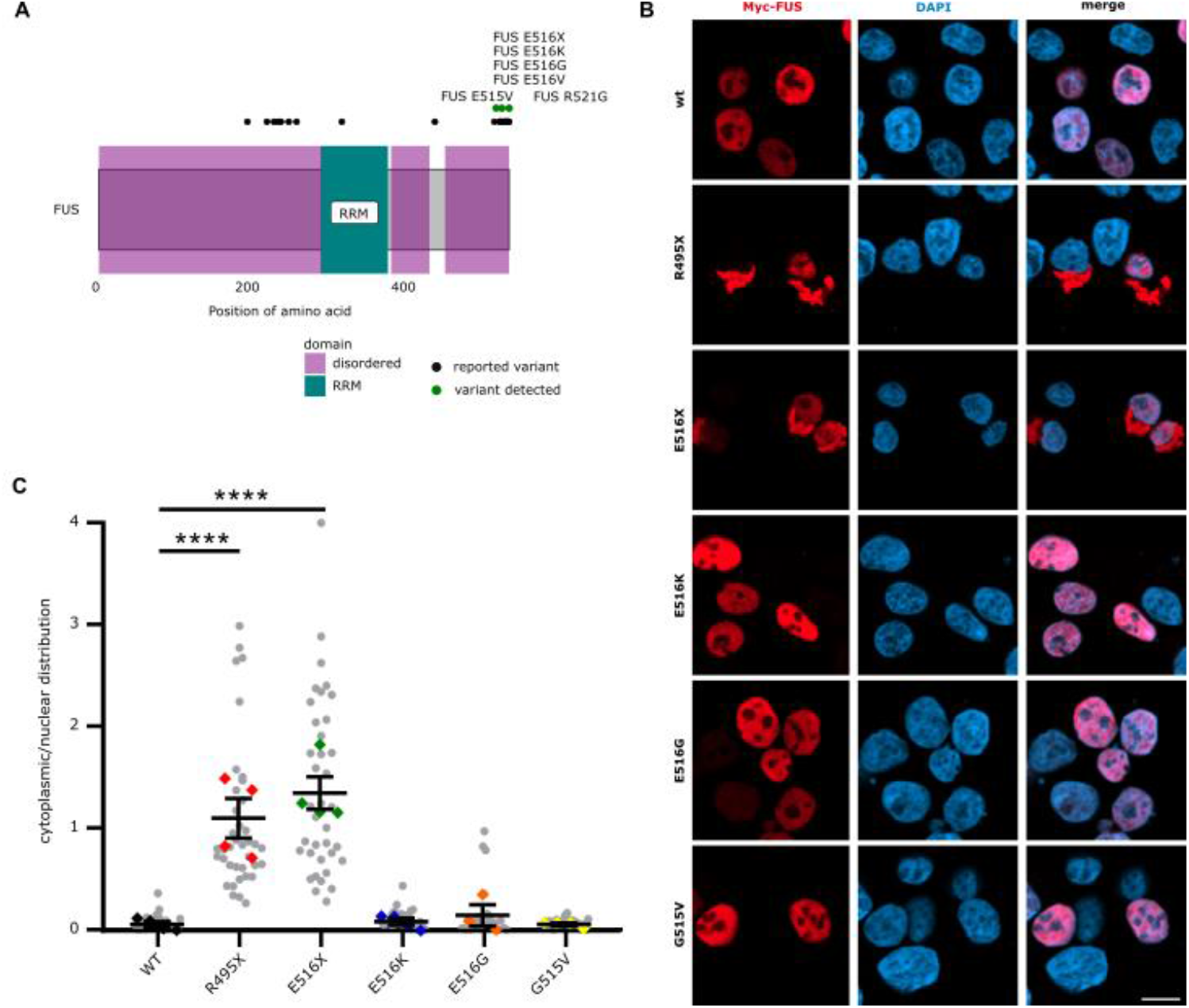
Functional validation of variants in FUS observed as somatic mosaic variants in sALS patients. (**A**) FUS protein structure with functional domains and known SNVs and SNVs detected in this study. (**B**) Cellular localization of wild type and mutant FUS in cultured HEK293 cells. Scale bar 10 μm. (**C**) Ratio of cytoplasmic to nuclear localization of FUS protein for the different FUS variants.

### Increased somatic variant burden in excitatory neurons identified by variant calling from single cell sequencing data

To further understand the cell type-specific distribution of somatic mosaic variants in the motor cortex, we performed somatic variant calling in single cell RNA-sequencing data from a publicly available cohort of sporadic and familial ALS (sALS and fALS), sporadic and familial FTLD (sFTLD and fFTLD), and control samples using Scomatic (Fig. 3, Supplementary Table 5)^18^. In total, we processed around 850,000 cells with their original cell type annotation.

**Figure 3.**
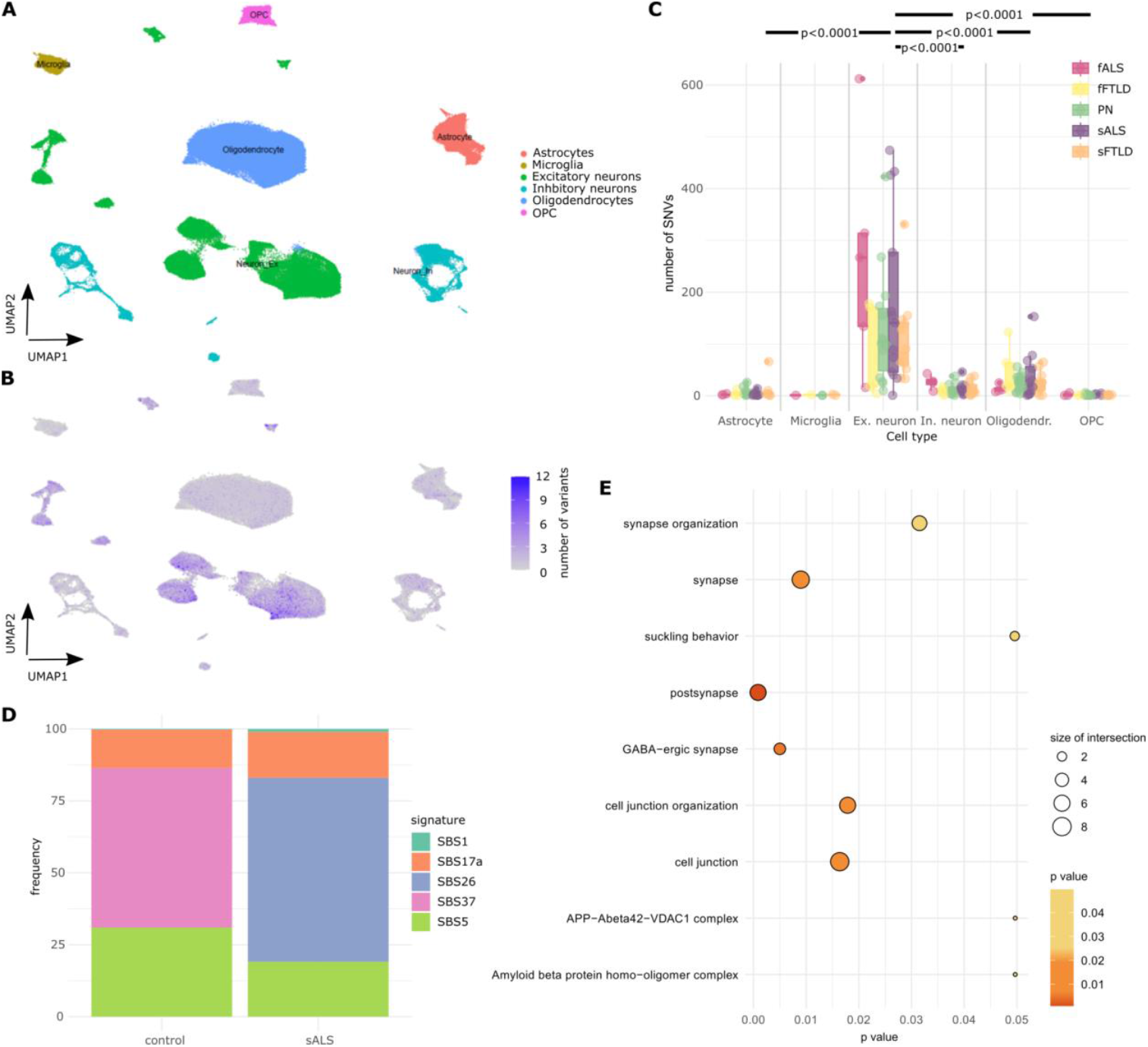
Somatic variant calling from single cell RNA-sequencing data. (**A**) Dimension reduction and visualization of single cell RNA expression levels of clusters of cells by Uniform Manifold Approximation and Projection (UMAP) of a dataset with sALS, sFTLD, and annotation of cell types. (**B**) UMAP of cell populations with the color indicating the number of all (coding and non-coding) somatic variants detected by SComatic per cell. (**C**) Distribution of number of variants per cell type and disease status. Pairwise statistical analysis was performed by using a generalized linear model corrected for multiple testing and number of cells and shown exemplarily for sALS samples. (**D**) Relative fraction of DNA damage-related single base substitution signatures in controls in comparison to sALS samples. (**E**) Gene sets enriched for SNVs in excitatory neurons.

Our analysis revealed that excitatory neurons were the cell type with the highest mutational burden for diverse ALS diagnoses (Fig. 3A-C). Two clusters of subpopulations of excitatory neurons were identified, in which the numbers of SNVs were especially high (clusters 5 and 9, Supplementary Figure S6). These clusters were mainly composed of sALS cells, revealing a selective high vulnerability of these cells to accumulate SNVs.

By using all somatic variants detected in the excitatory neurons, we computed *de novo* mutational signature profiles per condition. We observed a trend towards an enrichment of DNA damage signatures, e.g. SBS26 resulting from defective DNA mismatch repair, in sALS patients compared to controls (Fig. 3D). Gene set enrichment analysis showed an increase in somatic SNV burden in genes related to synapse and cell junction organization in sALS patients (Fig. 3E).

Taken together, our analysis identified an enrichment of somatic mutations predominantly in excitatory neurons, with an association to DNA damage repair signatures and cell junction and synapse organization, suggesting that altered DNA repair mechanisms may contribute to sALS pathogenesis.

## Discussion

Here, we show that sporadic, but not monogenic ALS caused by germline mutations, is linked to somatic mutations in the motor cortex. We also demonstrate ALS-typical pathology of FUS protein harboring a specific mosaic mutation in sALS patients. Respective mutations usually disturb the nuclear localization sequence of the protein, leading to nucleocytoplasmic redistribution and cytoplasmic aggregation^22^, accompanied by nuclear loss-of-function effects^23^. We observed a robust cytoplasmic mislocalization and deposition of the FUS p.E516X mutant protein found in the mosaic state (7% allele frequency in an sALS patient) in cultured cells. It is important to emphasize that, if found as a germline mutation, this would have been classified as ALS-causative.

Moreover, analysis of single cell transcriptomic data revealed an increased burden of somatic mutations in human motor cortex neurons, supporting a neuronal origin for sALS and that neurons are more prone to accumulate somatic mutations than other brain cell types^24^. Excitatory neurons show an even higher enrichment of mosaic mutations than inhibitory neurons, in line with the view that ALS starts in excitatory neurons^25^. The data support a pathogenic role of low-frequency, somatic mutations in sALS patients.

Notably, in several sALS motor cortices, we detected somatic variants in more than one ALS gene, suggesting a poly- or oligogenic mosaic origin. It remains to be shown whether the different variants detected in an individual arose from the same cells or in different cells in a “colony” or even in cell types that could communicate to instigate pathology. The age-related accumulation of variants in neurons and genomic instability as an effect of defective DNA damage response should be considered as a sALS pathomechanism^26,27^. Many of the known ALS-related genes impair DNA damage repair directly or indirectly^28^. Thus, an accumulation of variants in DDR-related SBS signatures as observed for excitatory neurons might contribute to the development of sALS. In summary, this study sheds new light on the origin of sALS. In perspective, these findings may have important implications for therapeutic design, because the identification of mosaic mutations could render respective patients suitable for gene-specific interventions, based on already successfully adopted antisense-oligonucleotides and siRNAs^29^. A necessary prerequisite will be the development of sensitive procedures for the diagnosis of ALS and for the reliable identification of mosaic mutations in biofluids from ALS patients.

## Acknowledgements

The authors would like to thank the patients and their families. We acknowledge the support by the Sequencing Core Facility of the German Cancer Research Center and the Omics IT and Data Management Core Facility with special thanks to F. Petermann, M. Vogel, L. Weiser, and G. Warsow. We acknowledge the support of the LIMa Live Cell Imaging Mannheim at Microscopy Core Facility Platform Mannheim (CFPM). We thank K. Hauschulz for suggestions on sequencing library preparation. LV wishes to thank N. Paramasivam and S. Uhrig for discussions on variant calling. The project was in part funded by a grant by the AI Health Innovation Cluster to RP, BB, LV, and JW (AIH23). R. Yilmaz was supported by the Deutsche Forschungsgemeinschaft Walter Benjamin Programme (YI 209/1-1, AOBJ 680080).

## Author contributions

OGV analyzed the variants from targeted sequencing and single-cell datasets, prepared graphs, and wrote the manuscript. RP designed the project, prepared patient material for sequencing and performed experimental validation in patient material, prepared graphs, and wrote the manuscript. RY designed, established and performed sequencing validation of variants. AF, LD, and SM performed experimental validation of variants in cultured cells. XY analyzed data. NT contributed performing targeted amplicon sequencing. DB, JCMS, and BB supervised aspects of the project. PMA and KMEF prepared and provided patient material. LV designed and supervised the project, established the analysis pipeline for variant calling from targeted sequencing, prepared material for sequencing, analyzed data, prepared figures, and wrote the manuscript. JW designed and supervised the project and wrote the manuscript. All authors have revised or critically reviewed the article.

