## Supplementary Figure for "Somatic mutations in ALS genes in the motor cortex of sporadic ALS patients"

# -

### Supplementary Text

##### Disease-related germline variants detected in sporadic ALS patients

We first identified heterozygous germline mutations from the targeted sequencing approach. From this analysis, we detected three non-synonymous, heterozygous germline SNVs in two patients diagnosed with sALS. None of these SNVs were detected in any of the controls. One of the SNVs KIF5A p.A268T, detected in patient 20 and reported to have a pathogenic effect at the clinical level, has previously been linked to spastic paraplegia (dbSNP[1]: rs139015012). We observed that the patient did not show any additional somatic SNVs in any of the known ALS-related genes of the targeted panel. Another sample (patient 19) showed a heterozygous mutation in ALS2 p.T1472M (rs201089588), which has been reported previously in two ALS cases and one case of infantile-onset ascending hereditary spastic paralysis[1]. The same patient carried a second heterozygous germline variant in SQSTM1 p.P387L, which was reported as “likely pathogenic” in ClinVar (rs776749939) with studies linking this variant to increased susceptibility to frontotemporal dementia (FTD) and ALS[2,3]. Therefore, we excluded patients 19 and 20 from any quantitative analysis, because they carried germline mutations that were likely pathogenic and not detected by previous examination.

##### Mutational signatures enriched in sALS patients

The distribution of base exchanges within its trinucleotide context observed by deep targeted sequencing in sALS patients and controls showed a similar pattern with most base exchanges occurring as C>A, C>T, T>A, T>C, and T>G (Supplementary Figure S5A-B). Base exchange frequencies for C>G were significantly lower for both, controls and sALS samples.

We analyzed single base substitution (SBS) signatures and compared them between the groups of samples. A number of single base substitution signatures can be related to underlying mechanisms for variant accumulation in the genome. We observed an enrichment of the signatures SBS2, SBS85, and SBS89, related to APOBEC and cytidine deaminase, in sALS samples compared to controls (Supplementary Figure S5C). Furthermore, the signatures SBS8, SBS10b, and SBS11, although appearing with a small contribution, were only detected in sALS samples. These signatures are associated with unknown aetiology of CC > AA exchanges, polymerase epsilon exonuclease domain mutations, and the exposure to alkylating agents, respectively[4]. We observed an increase in SNVs as part of the signature SBS89 of unknown aetiology for sALS patients as well as a decrease in signatures SBS29 and SBS25 potentially related to tobacco and chemotherapy (Supplementary Figure S5D). Furthermore, we observed an overall higher cosine similarity with mismatch repair deficiency signatures for sALS patients compared to controls (Supplementary Figure S5E). This supports a potential role of DNA damage repair deficiency in sALS development.

**Somatic mosaic variants detected in sALS patients**

We compared the occurrence of somatic variants at an AF between 1% and 35% in genes known to be involved in ALS between fALS, sALS, and control samples and performed validation of the variants by amplicon sequencing (Supplementary Figure S7, Supplementary Table 6). In controls, we detected somatic variants in *FIG4*, *GLE1*, *TUBA4A*, *TBK1*, *SETX*, and *FUS*, as well as an SNV in a 3'UTR of *VAPB*. fALS patients additionally showed variants in *CCNF*. For sALS patients, we observed a significant increase in the patients carrying variants in *FIG4* (9/9), *GLE1* (9/9), and *TBK1* (8/9). Variants in SETX were observed in 6 of 9 and variants in *FUS* in 4 of 9 sALS cases. Additionally, we observed exonic non-synonymous variants in *ALS2*, *NEFH*, as well as SNVs in the 5'UTR of *VAPS* in 44% of cases and in 3'UTRs of *NEK1*, *PFN1*, and *TARDBP* unique to individual patients. In comparison, we only detected 2 SNVs in 3'UTRs in *TARDBP* in fALS patients and none in controls, which occur in the general European population at a frequency of 3.2% and 0.13%.

Mutations in *TARDBP* can cause ALS by a loss-of-function principle, accompanied by cytoplasmic aggregation and loss of nuclear TDP-43 at the protein level. To assess the potential functional importance of the 3'UTR variant on *TARDBP* expression, we investigated evolutionary conservation using multiple sequence alignment across a wide range of species (Supplementary Figure S8). We focused on the sequence region of the 3'UTR variants in *TARDBP*. Results revealed a 100% sequence conservation across species analyzed in the region harboring the somatic mosaic low AF variant. Evolutionary conservation was less pronounced in the two regions, for which two fALS patients carried heterozygous variants (with 17 or 28 and 22 of 28 species showing variability in each of the regions, respectively). The evolutionary conservation between mosaic sALS and heterozygous fALS 3'UTR variants in *TARDBP* might indicate the functional importance of the 3´UTR region in which the rare low AF variant from the sALS patient is located. It has been reported that TDP-43 is able to autoregulate its expression levels and localization. This autoregulation of TDP43 levels and localization is achieved in part via a negative feedback loop by self-targeting the TDP-43 binding region located in the 3´UTR of its transcript[5–7]. A previous study has reported a similar *TARDBP* 3'UTR variant in a frontotemporal lobar degeneration patient (FTLD) and a frontotemporal lobar degeneration patient with ALS co-morbidity (FTLD-ALS), both with TDP-43 proteinopathy[8]. The *TARDBP* 3'UTR variant was accompanied by a two-fold increase in total

*TARDBP* mRNA when compared to controls.

We observed multiple variants in *FIG4* (Supplementary Figure S7). Selection based on pathogenicity prediction by AlphaMissense or description of clinical effects in the literature retained one variant that was described to be pathogenic. This missense variant in FIG4 at amino acid position 157 substituting a valine by a leucine (FIG4 p.V157L; NM_014845:exon5:c.G469T) was found in sALS patient 30 with an allele frequency of 1.6%. This position corresponds to the start site of the phosphatase domain of FIG4. A variant at this exact same position but with a valine to methionine substitution (FIG4 p.V157M) was reported in the ClinVar database (rs1455052760), and the phenotype reported is linked to Charcot-Marie-Tooth disease type 4, a neurodegenerative disease that involves also motoneurons.

Furthermore, a non-synonymous pathogenic variant in the *SETX* gene (SETX p.A2127D) was found with an allele frequency of 1.2% in patient 28.

One of the sALS patients (patient 27) carried a high impact stop-gain variant in the *TBK1* gene (TBK1 p.Q655X) with an allele frequency of 3.9% (Supplementary Table 4). However, the same TBK1 p.Q655X variant was also detected in a control sample (patient 18) with a slightly lower allele frequency than 1.5%, suggesting that additional factors contribute to disease development.

We observed an enrichment of *FUS* variants in exon 15 in sALS cases (Supplementary Figure S7). Exon 15, the extreme C-terminus of the protein, encodes the nuclear localization signal (NLS) and the glycine-rich domain of the protein that enhances the aggregation-prone property of FUS protein[9,10]. It interacts with the nuclear import receptor transportin (karyopherin β2), essential for shuttling FUS between the nucleus and the cytoplasm[11]. Mutations in the NLS domain of FUS lead to cytoplasmic translocation and aggregation of FUS and cause ALS[12–14]. Here, we report six distinct low AF missense somatic mutations within the NLS domain of FUS, one stop-gain (FUS p.E516X), and four non-synonymous SNVs (FUS p.E516K, p.E516G, p.E516V, and p.R521G), all found in sALS patients. As a first step, we used AlphaMissense to assess the functional impact of these mutations. All of the variants were flagged as “pathogenic”. Only one control reported a missense somatic variant in this very same FUS region (FUS p.G515V), which was the only variant predicted as benign by AlphaMissense. FUS p.R521G missense mutations in FUS have been previously reported in a study of three different fALS pedigrees. The variant led to a cytoplasmic retention of the mutant protein observed both in brain tissue and transfected N2A and SKNAS cells[12]. Additionally, we identified a somatic mutation in three sALS patients reporting an adenine to cytosine substitution in the acceptor splice site of intron 14 (NM_004960:exon15:c.1542-

2A>T). This specific splice-site mutation (rs1266751712) was identified as the most probable cause of ALS in a screening of a large French family[15]. Furthermore, the authors showed that the mutant allele was missing the 40 bp coding sequence of FUS exon 15 as well as the first 203 bp of the 3′ untranslated region.

Moreover, our results showed that three sALS patients carried somatic variants of low allele frequency and of functional importance in at least two positions in the same or different ALS-related genes. For patient 28, we detected the variants FUS p.E516V, FUS p.R521G, SETX p.A2127D, as well as mutations in canonical FUS and TARDBP splice sites. In patient 19, we observed FUS p.E516G and FUS splice site variants (c.A1542-2T), and in patient 24, FUS p.E516K and another FUS splice site variant (c.A1542-2T) were observed.

Taken together, we identified several pathogenic somatic mutations in ALS-associated genes, suggesting somatic mosaic variants to play a role in sALS development.

**Detection of variants in fALS-related genes and beyond from single cell RNA-sequencing**

We investigated our panel of known fALS-related genes by selecting all variants identified from single cell RNA-sequencing data, even if only detected in a single cell type. We observed a total of 15 variants in the selected genes in excitatory neurons (10 intronic and 5 UTR3). Overall, the genes with the largest average number of variants per patient were *MAPT*, *KIF5A*, *NEK1*, *ALS2*, and *FIG4*.

We also investigated somatic variants in other genes that were detected in excitatory neurons comparing multiple cell types (presence of a variant in a specific cell-type but not in the others). We detected genes with an increased mutational burden in sALS by computing the ratio of the number of variants between sALS samples and controls for each of the genes, selecting genes that showed a log2-difference larger than 2 and not showing any SNVs in the control samples (Supplementary Figure S6E, Supplementary Table 7). These variants were mostly observed in genes with potential functional roles in brain development and neurodegenerative diseases. Gene-set enrichment analysis of the top mutated genes, with a log2-difference larger than 2, related them to synapse, GABA-ergic synapse, synapse organization and postsynaptic functions (Fig. 3E).

### Supplementary Materials and Methods

#### Patient cohort

Fresh frozen autoptic human precentral gyrus (and spinal cord) tissues of donors with ALS and control donors (with similar age and sex distribution) were provided by the ALS Brain Bank at Umeå University in Sweden and the Netherlands Brain Bank. The Swedish ALS patients were diagnosed in accordance with the EFNS Guidelines for the management of ALS[16]. Using genomic DNA, all patients were Sanger sequenced for a panel of known fALS-causing genes. Patients with a pathogenic mutation in one of these genes or patients with a definite family history of ALS within three generations were termed fALS, all others were termed sALS. The autopsies were performed prospectively since 1993 adhering to a specific protocol and the tissue specimens saved in small pieces at -80°C until use. Control samples were collected and stored by using the same procedure. A complete list of samples used for targeted sequencing is found in Supplementary Table 1.

#### DNA extraction and library preparation

Genomic DNA was extracted from ca. 20 mg of tissue by the QIAamp DNA Mini kit (QIAGEN, Germantown, USA) according to the manufacturer’s recommendations. For whole exome sequencing the DNA was further processed by CeGaT Gmbh (Tübingen, Germany). For targeted sequencing, 50 ng of genomic DNA was used for library preparation and target enrichment by the SureSelect XT HS2 DNA system (Agilent Technologies, Santa Clara, CA, USA).

#### Design of panel for target amplification

A DNA panel for Agilent SureSelect XT Library Preparation was designed targeting the exonic and adjacent intronic regions of a total of 28 genes covering a genomic region of 1012 kB (Supplementary Table 2). 3358 probes with a total probe size of 107 kbp were designed by using the Agilent SureDesign software covering the coding exons of the genes and extending into the 3’ and 5’ end by ~50 bases each. Details about the target ID, intervals, and regions are provided in Supplementary Table 2.

#### Target enrichment and sequencing library preparation

The Agilent SureSelect XT HS2 DNA Library Preparation and Target Enrichment protocol was followed for target enrichment. All steps were performed as described in the protocol provided by the manufacturer with the following adjustments: 23-50 ng of genomic DNA were used for enzymatic fragmentation for 15 min to obtain a target fragment size of 150-200 bp. During the pre-capture PCR 9-10 amplification rounds (for 50 ng and for 23 ng of input DNA, respectively) were performed. The quality of the libraries was followed by using the D1000 ScreenTape System (TapeStation System, Agilent).

**Targeted next-generation DNA sequencing**

Sequencing of a pool of 16 samples each with a concentration of 10 nM each was performed at the Sequencing Core Facility of the German Cancer Research Center. The samples were sequenced on a HiSeq 4000 instrument (Illumina) by paired-end sequencing of 100 bp in a 4-color patterned flow cell with 235 M reads targeting a coverage of ≥2000x. Per sample, ~10-20 Mio reads were obtained.

**Targeted DNA sequencing analysis**

A schematic overview of the analysis workflow is provided in Figure S1. Reads from individual samples in raw FASTQ files were demultiplexed based on the Agilent indices used. Quality control of sequencing results was performed by using FastQC and MultiQC[17,18]. The molecular barcode sequences were extracted and sequencing adaptors were removed from the raw reads by using AGeNT Trimmer v2.0.3. Reads were then mapped to the reference genome hsGRCh37d5 by using BWA-MEM[19] v0.7.17. The molecular barcode tags were added to the aligned reads in the BAM file by using SAMtools view. Error-corrected consensus reads were generated by using the molecular barcodes and AGeNT LocatIt v2.0.5 in v2Duplex mode with a barcode distance of 0. BAM files were validated by using picard v.2.25.1 ValidateSamFile, read groups were added by SAMtools v1.12 addreplacerg[20], if required, and BAM files were sorted and indexed (SAMtools). Variant calling was performed by multiple variant callers. Octopus v0.7.4[21] was run in the *cancer* setting calling germline and somatic variants with random forest filtering and somatic forest filtering. SAMtools v1.12 mpileup was run with default settings and varscan v2.3.8 mpileup2indel and mpileup2snp were used with a minimum read depth of 8, minimum number of supporting reads of 2, minimum base quality of 15, and minimum variant allele frequency of 0.01. Variants had to be supported by ≤90% of the reads by one strand to be considered for downstream analysis. GATK v4.2.0.0[22] Mutect2[23] and Mutect2-PoN (GenomicsDBIImport, CreateSomaticPanelOfNormals) were run, the latter with a panel of normals consisting of nine control samples (see Supplementary Table 1). Variant calling by using GATK v4.2.0.0 HaplotypeCaller[24] was performed with the *ploidy* parameter set to 100 to consider all variants as different haplotypes. Strelka v2.9.5[25] was run in default mode and blat v3.6 MosaicHunter[26] was run in the exome and the genome modi. The variants detected in all tools were consolidated and DeepMosaic v1.0.0[27] *draw* and *predict* was used to detect somatic mosaic variants with model type efficientnet-b4.

##### Variant annotation, filtering and pathogenicity prediction

Variants were annotated using the annovar tool for variant effect prediction, and selected only if having a gnomAD frequency in the general population of < 0.001[28,29].

Predictions for all possible single amino acid substitutions were obtained from AlphaMissense[30] (AlphaMissense_aa_substitutions.tsv.gz file from https://zenodo.org/records/8208688, 2023-09-19). The database of predicted effects was split by gene, and individual coding mutations found on the panel that have passed all filters were interrogated.

##### Selection of range of allele frequencies

For selecting the allelic frequency (AF) range in which mosaic variants could be identified reliably, we included control and spike-in DNA samples from a patient with the SOD1 p.D90A homozygous germline variant at known concentrations (1.5% and 0.5%). After applying our pipeline for variant calling, we identified somatic variants down to a spike-in variant AF of ~1%. Variants with AF below this cut-off, e.g. at 0.5% could not be detected by any of the variant callers used in their default configurations, although the variant was visible at the expected base exchange frequency in genome browsers. Therefore, we determined a lower threshold for detecting somatic mosaic variants at a variant allele frequency of 1%, and focused the analysis on variants occurring at an allele frequency between 1% and 35%.

##### Statistical analysis of mutation frequency

Mutation enrichment test between disease categories were analyzed using negative binomial generalized linear models (GLM) to account for overdispersion in count data. The model included disease category (sALS, fALS, control) as the main predictor, with sex, age, origin of patient, and total coverage as covariates to control for potential confounding effects.

Model fitting was performed using the glm.nb() function from the MASS package in R. Test for disease * sex interactions was done using likelihood ratio tests to determine whether category effects varied by sex. Since no significant interaction was detected ($p = 0.285$), we proceeded with the additive GLM model.

Pairwise comparisons between disease categories were conducted using estimated marginal means with FDR correction for multiple testing, implemented via the emmeans package. Statistical significance was set at α = 0.05.

**Validation and quantification of mosaic mutations with targeted amplicon sequencing**

Targeted amplicon sequencing was performed for candidate variants detected from deep panel sequencing as described above, to independently validate the identified mosaic mutations. PCR products were designed with a target length between 170 and 190 bp. PCRs were performed using the GoTAq Colorless Master Mix (Promega, M7832) on the ALS samples and controls as reported previously[31]. The libraries were sequenced on the Illumina NextSeq 550 platform with 151 bp paired-end reads using the NextSeq 500/550 Mid Output kit (300 cycles). The analysis was performed by comparing binomial distributions as described in Chung et al. (2023)[31]. A full validation was achieved, if the difference in the binomial distribution was significant against three control samples, whereas a partial validation was achieved when the difference was significant against at least one of the control samples.

**Single cell RNA-sequencing analysis and somatic variant calling**

We downloaded the raw fastq files from a public single cell RNA-sequencing dataset of sALS, fALS and controls, and additionally including sFTLD and fFTLD[32] from NCBI. In total we processed around 850.000 cells (n. of samples: c9ALS=5, c9FTLD=5, control=15, sALS=13, sFTLD=12). Fastq files were processed using cellranger version 8.0.1, and human genome reference version GRCh38. We then used SComatic for somatic variant calling analysis following the guidelines and scripts provided by the authors[33], annotations of groups of cells for SComatic were defined using the data´s original authors annotation of cell types, namely, astrocytes, microglia, excitatory neurons, inhibitory neurons, oligodendrocytes, and oligodendrocyte precursors (OPCs). Bedtools version 2.29.2, and annovar for variant annotation were used as part of the variant calling pipeline. Variants were filtered by selecting the ones with the PASS flag, as described in the SComatic documentation. Downstream analysis was carried out in R version 4.3.0, using Seurat version 5.1.0 and SeuratObject 5.0.2[34]. We selected the top 2000 most variable genes using FindVariableFeatures, and computed PCA and UMAP with those. Clustering was done as follows: FindNeighbors was computed using the first 25 PCs, and FindClusters with a resolution of 0.8. Finally, information of somatic variants per barcode was appended to the Seurat object metadata. Gene markers for clusters 5 and 9 were selected using FindMarkers specifying as ident.1 the cluster of interest. Gene enrichment and ontology analysis for the selected markers was done using the gprofiler2 package with correction method g_SCS, and using the top 500 markers for the mentioned clusters.

Computation of *de novo* mutational signatures and decomposing the *de novo* signatures into the

COSMIC signatures was done using the R package SigProfilerExtractorR version v1.1.16[35].

28. Wang K, Li M, Hakonarson H. ANNOVAR: functional annotation of genetic variants from high-

throughput sequencing data. *Nucleic Acids Research*. 2010;38(16):e164-e164. doi:10.1093/nar/gkq603

#### Supplementary Figure S1

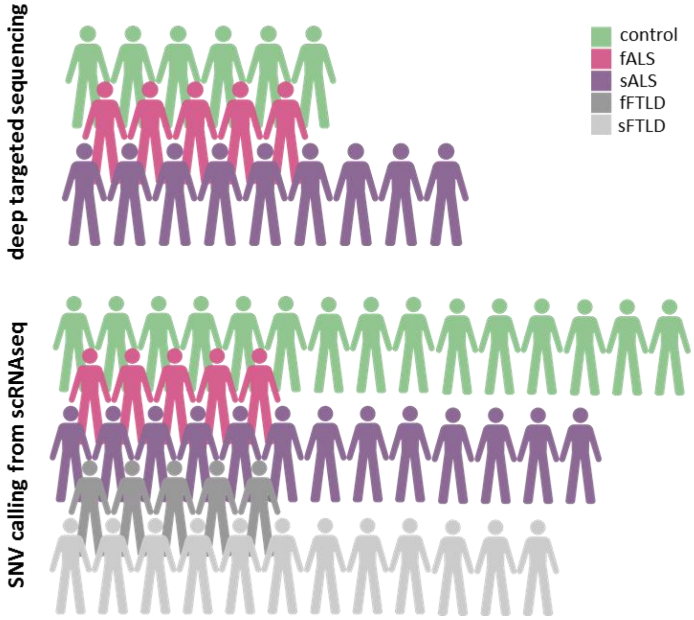


*Supplementary Figure S1. Schematic representation of the cohort of samples analyzed by deep targeted sequencing (upper panel) and by SNV calling from scRNAsequencing (lower panel).*

#### Supplementary Figure S2

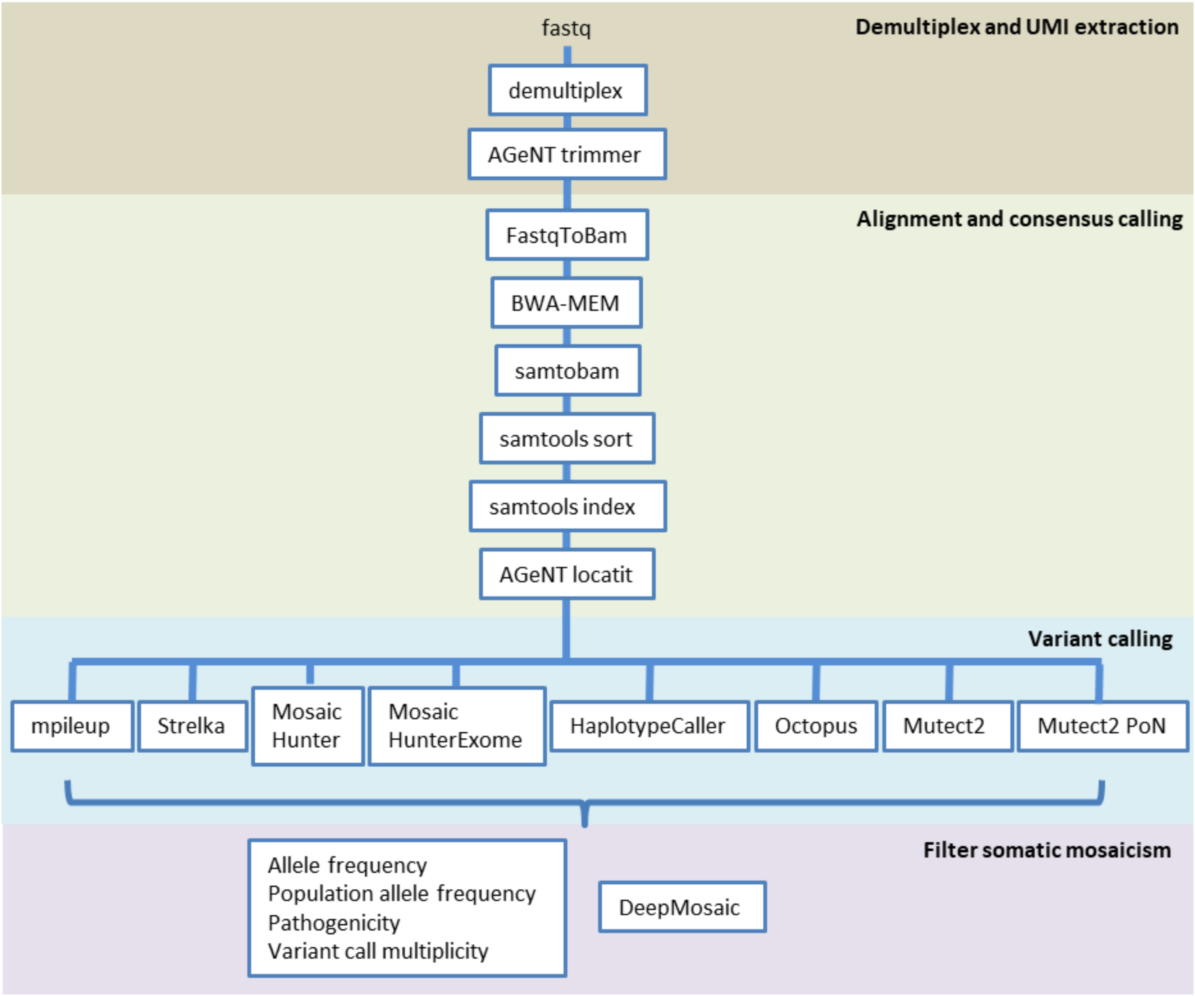


*Supplementary Figure S2. Schematic overview of the analysis approach for deep targeted sequencing data. After alignment and consensus calling based on unique molecular identifiers, variants were called by using multiple variant calling approaches and filtering of variants was performed by using DeepMosaic or a combination of cut-offs for allele frequency, population allele frequency, variant call multiplicity, and pathogenicity.*

#### Supplementary Figure S3

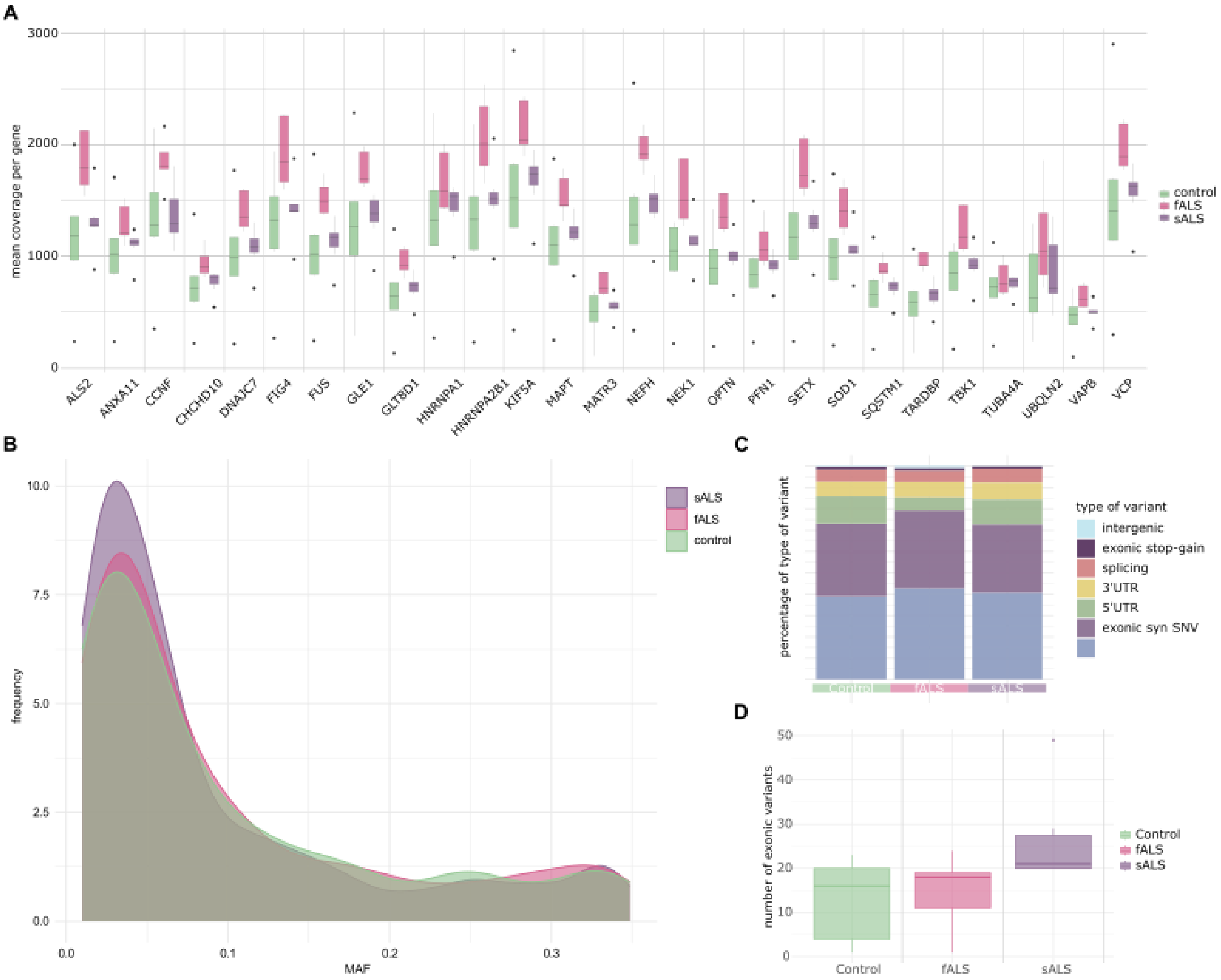


*Supplementary Figure S3. Comparative analysis of sequencing depth and quality. (A) Mean coverage per gene per disease condition. (B) Relative frequency of SNVs by allele frequency for each of the disease conditions. (C) Frequency of variant types detected per sample group. (D) Number of exonic variants detected per variant allele frequency for the different groups of samples.*

#### Supplementary Figure S4

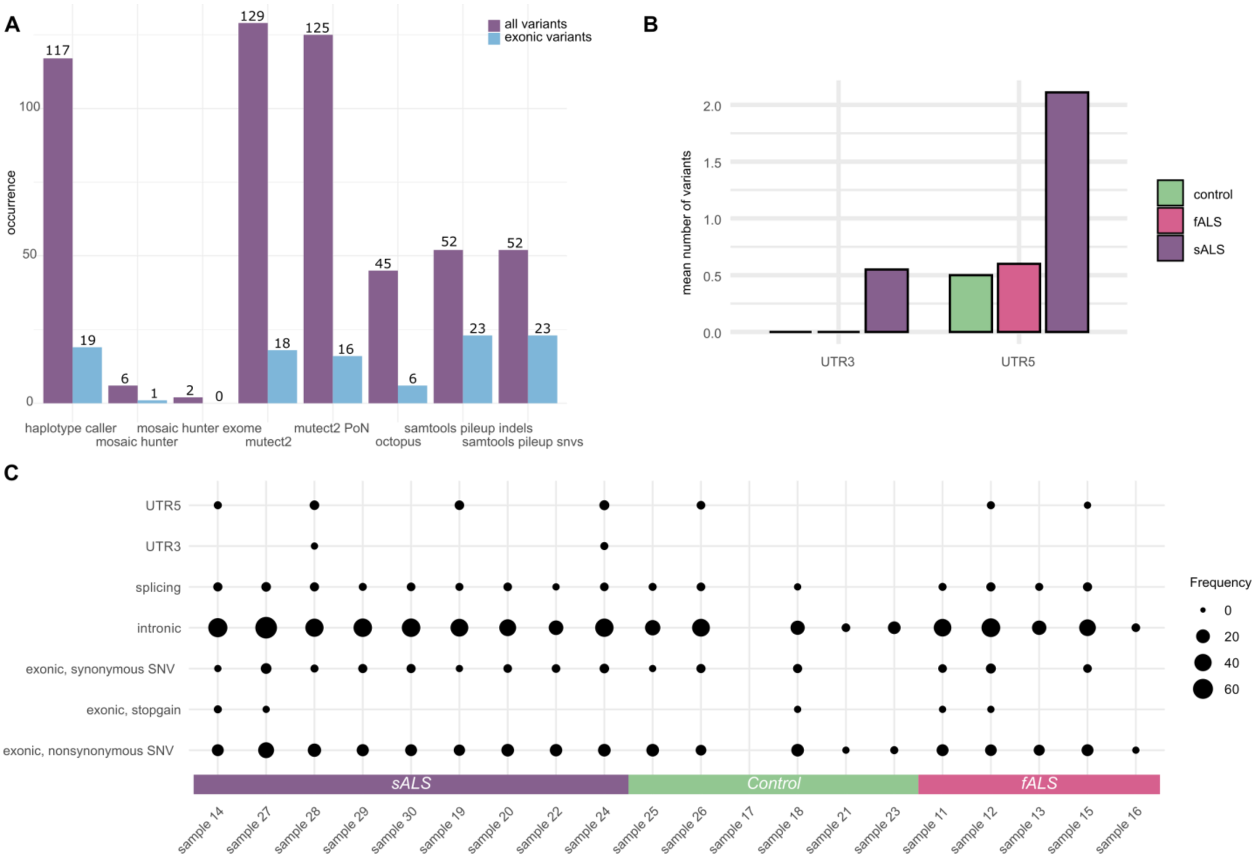


*Supplementary Figure S4. Single nucleotide variants from deep targeted sequencing. (A) Number of variants detected by using different variant calling approaches. (B) Mean number of variants detected in 3'UTR and 5'UTR regions in controls and ALS patients. (C) Frequency of variants in different regions per sample.*

#### Supplementary Figure S5

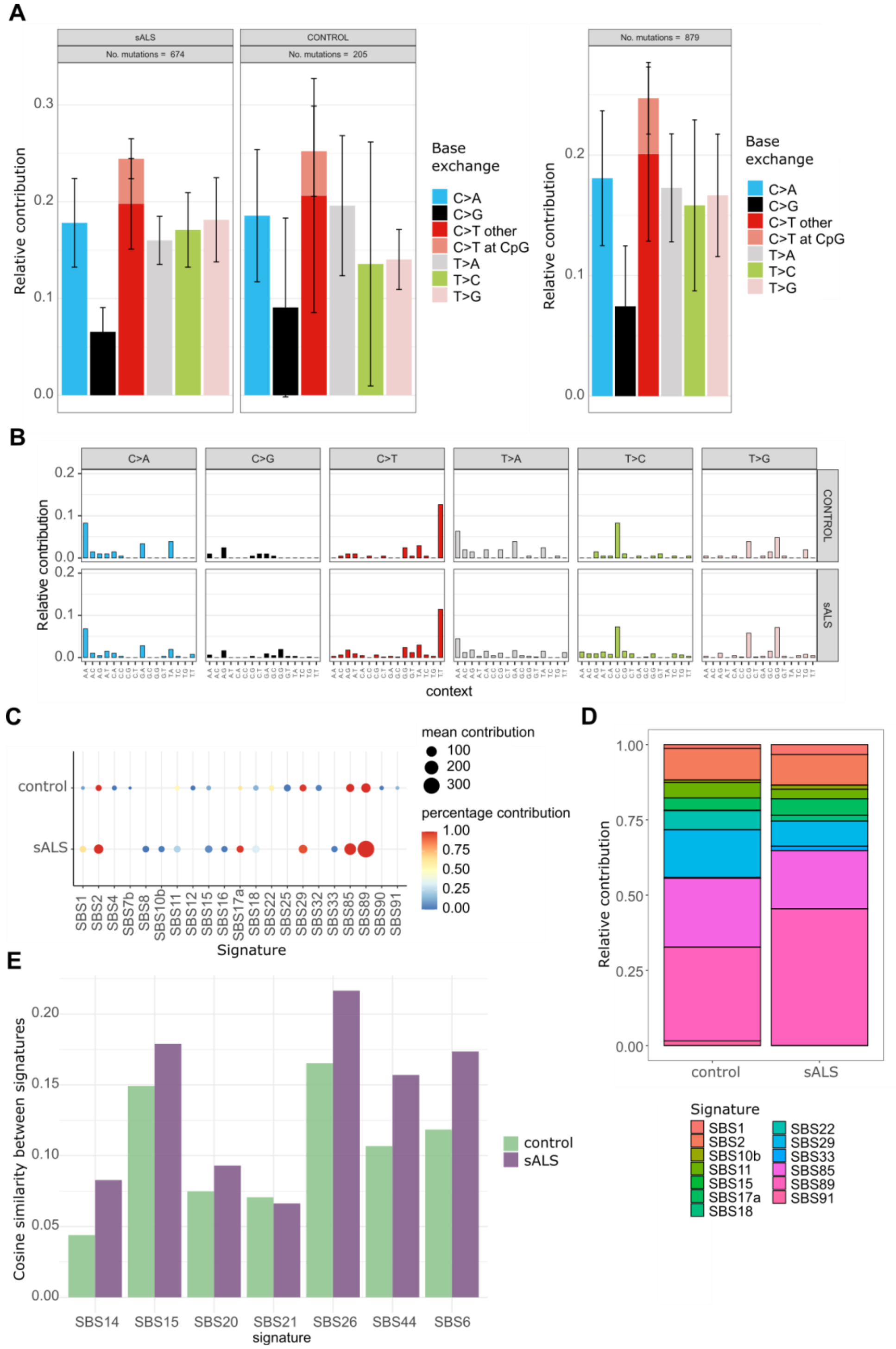


*Supplementary Figure S5. (A) Distribution of base substitutions in sALS samples and healthy controls. (B) Relative contribution of base substitutions in trinucleotide context between controls and sALS samples. (C) Single base substitution (SBS) signatures obtained from targeted panel sequencing of genes involved in ALS. Occurrence of SBS signatures in healthy individuals and sALS patients. (D) Relative contribution of single base substitution signatures to the signatures observed in healthy controls versus sALS patients. (E) Cosine similarity of signatures to mismatch repair-related signatures.*

#### Supplementary Figure S6

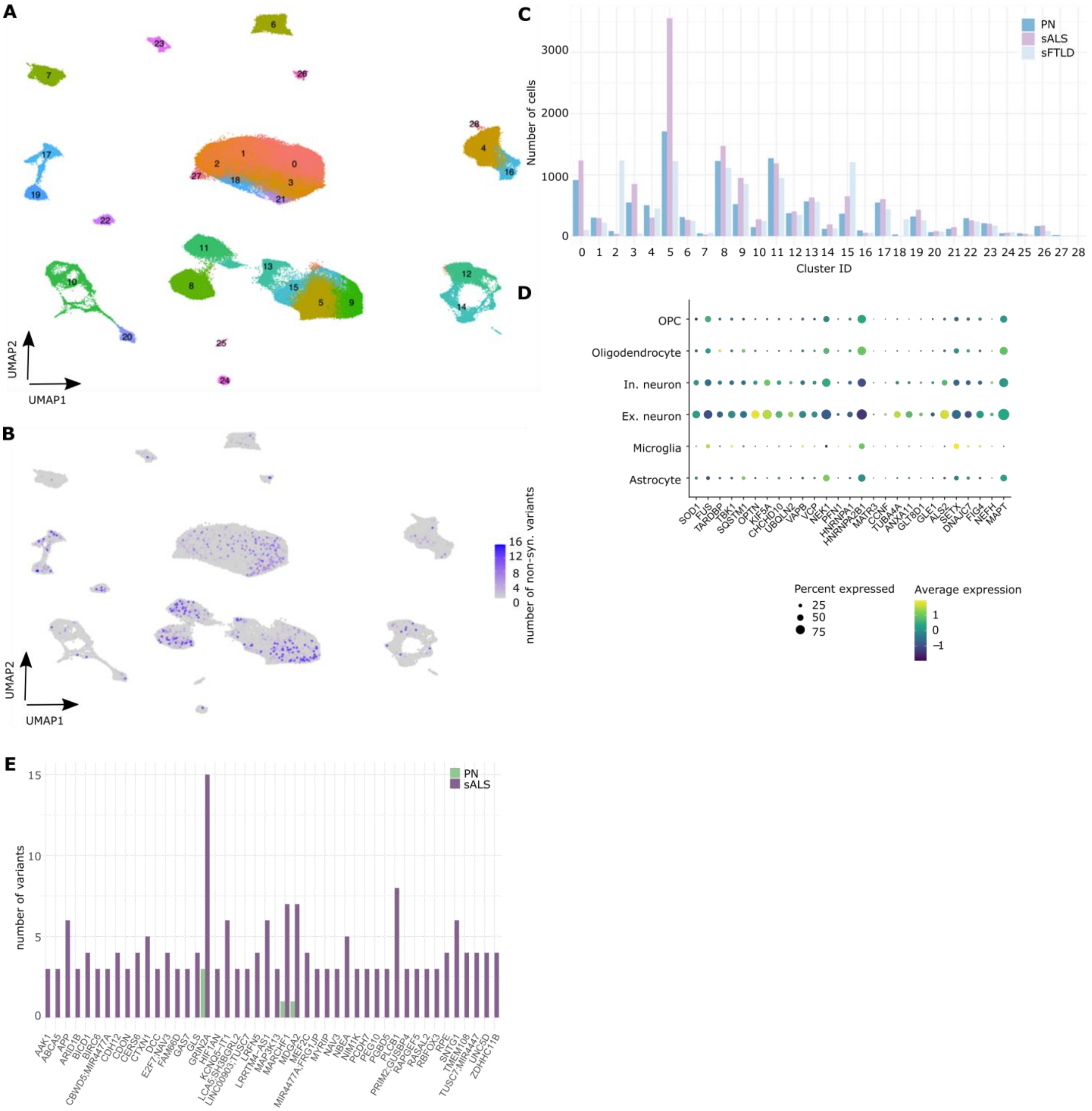


*Supplementary Figure S6. (A) Dimension reduction of single-cell RNA expression levels of clusters of cells by Uniform Manifold Approximation and Projection (UMAP) of a dataset with sALS, sFTLD, and annotation of cell types with unsupervised clustering. (B) UMAP of cell populations, color indicating the number of non-synonymous somatic variants per cell barcode.(C) Bar diagram showing the number of variants per cell type and disease status. (D) Gene expression of the genes included in the targeted deeply sequenced panel. (E)Number of single nucleotide variants per gene in excitatory neurons for controls and sALS samples.*

### Supplementary Figure S7

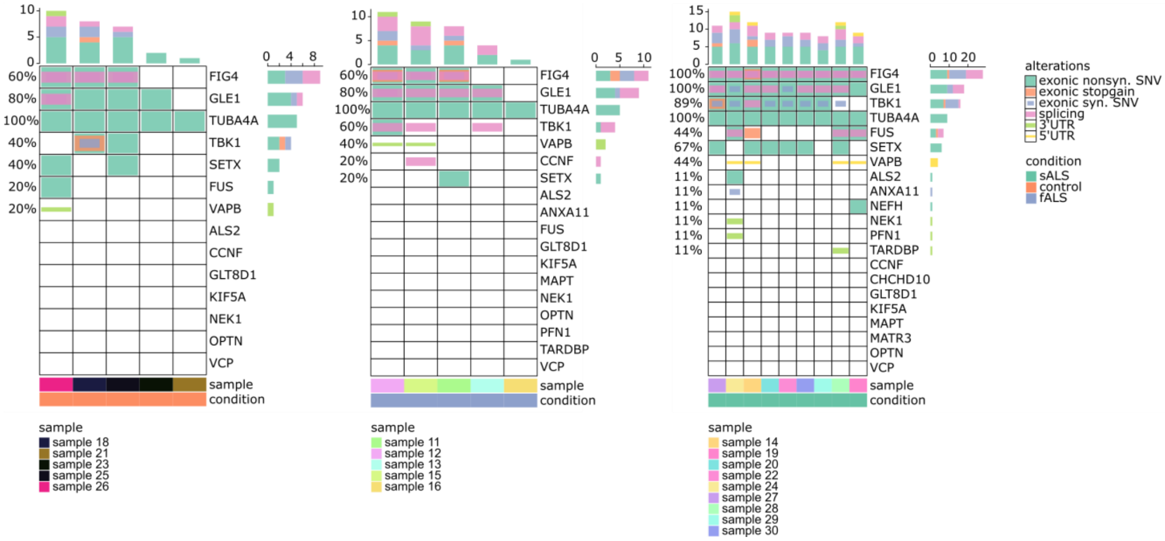


*Supplementary Figure S7. Frequency of somatic alterations in fALS genes of interest in control (left), fALS (center), and sALS (right) cases.*

### Supplementary Figure S8

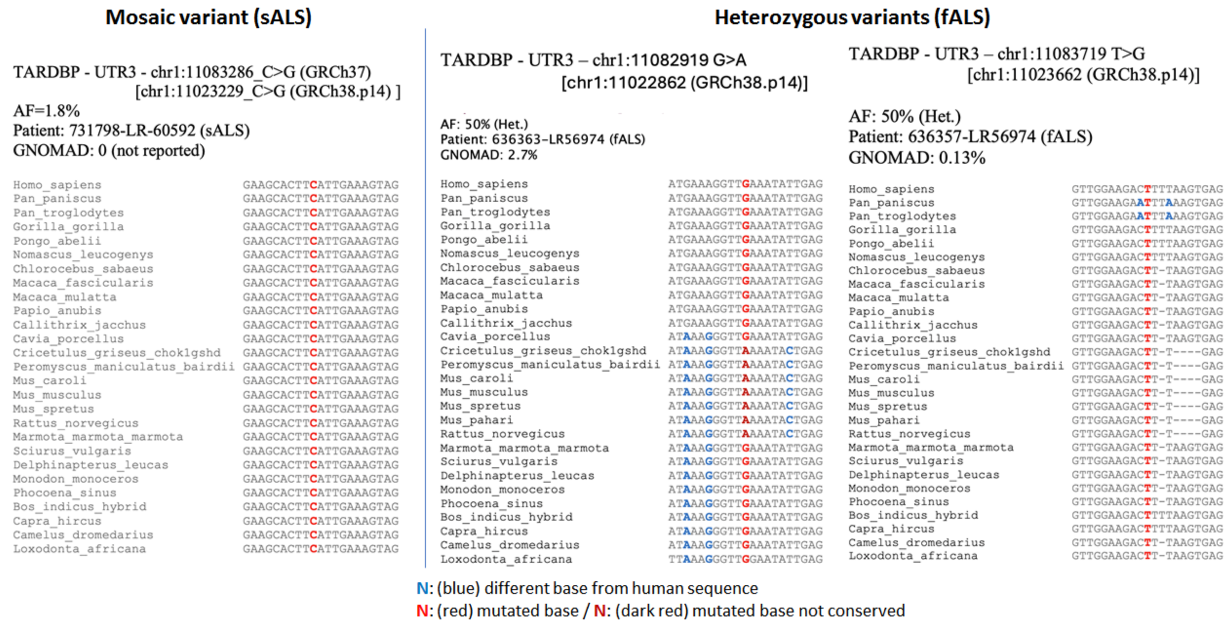


*Supplementary Figure S8. Evolutionary sequence conservation of TARDBP 3'UTR region in comparison for mosaic variants in sALS and heterozygous variants in fALS samples.*
